# PHluorin-conjugated secondary nanobodies as a tool for measuring synaptic vesicle exo- and endocytosis

**DOI:** 10.1101/2024.09.23.614406

**Authors:** Svilen V. Georgiev, Silvio O. Rizzoli

## Abstract

Neuronal communication relies on synaptic vesicle recycling, which has long been investigated by live imaging approaches. Synapto-pHluorins, genetically encoded reporters that incorporate a pH-sensitive variant of GFP within the lumen of the synaptic vesicle, have been especially popular. However, they require genetic manipulation, implying that a tool combining their excellent reporter properties with the ease of use of classical immunolabeling would be desirable. We introduce this tool here, relying on primary antibodies against the luminal domain of synaptotagmin 1, decorated with secondary single-domain antibodies (nanobodies) carrying a pHluorin moiety. The application of the antibodies and nanobodies to cultured neurons results in labeling their recycling vesicles, without the need for any additional manipulations. The labeled vesicles respond to stimulation, in the expected fashion, and the pHluorin signals enable the quantification of both exo- and endocytosis. We conclude that pHluorin-conjugated secondary nanobodies are a convenient tool for the analysis of vesicle recycling.

## Introduction

The accurate release of neurotransmitters is essential for neuronal communication, a tightly regulated process that is composed of several mechanisms, including the exo- and endocytosis of transmitter-loaded vesicles. The vesicles are typically categorized into three distinct pools, the readily releasable, recycling and reserve vesicles (Rizzoli & Betz, 2005), with the former participating primarily in exo- and endocytosis, while the latter is less responsive to synaptic activity. The dynamics of these vesicles, collectively known as the synaptic vesicle cycle, have long been studied by a plethora of techniques, and especially by the use of fluorescent probes (Kavalali & Jorgensen, 2014). Among the most widely employed such tools are the FM dyes, lipophilic fluorophores that are loaded into synaptic vesicles during endocytosis (Betz & Bewick, 1992; Hoopmann et al., 2012), and the pHluorins, genetically-encoded fluorescent proteins fused to the luminal domains of synaptic vesicle proteins, which are quenched by the acidic pH of the vesicles, and thereby report both exo- and endocytosis (Atluri & Ryan, 2006; Z. Li et al., 2005; Miesenböck et al., 1998).

Despite their substantial contributions to the field, these tools are not without inherent limitations. FM dyes, for instance, have broad excitation and emission spectra, which can be disadvantageous for multi-color imaging experiments, where spectral overlap must be minimized. Additionally, FM dyes exhibit relatively low brightness and photostability, characteristics that are critical for the accurate live imaging of the synaptic vesicle cycle (An et al., 2022). The use of FM dyes also necessitates prolonged washing steps, to remove the surface-bound molecules, and thereby reveal the ones trapped in endocytosed vesicles, and to enable exocytosis measurements, in which the FM dye loss from vesicles is measured. The long washing steps may result in the unintended loss of vesicular labeling, due to spontaneous exocytosis. Moreover, FM dyes can interact with various receptor types and potentially influence vesicle exocytosis, thereby confounding experimental results (Gale et al., 2001; D. Li et al., 2009; Mazzone et al., 2006; Zhu & Stevens, 2008). Finally, the FM dyes need to be introduced into the vesicles by a first round of stimulation, which may influence all subsequent experimental steps.

In contrast, the genetically-encoded pHluorins are packaged into vesicles by the secretory system of the neurons, and thus remove the problem of dye uptake. Since the development of the first pH-sensitive GFP fused to synaptobrevin 2 (Miesenböck et al., 1998), researchers have successfully engineered various other pHluorin chimeras, using synaptotagmin (Dean et al., 2012), vesicular glutamate transporter 1 (vGlut1) (Voglmaier et al., 2006), synaptophysin (Granseth et al., 2006), or the vesicular GABA transporter (vGAT) (Santos et al., 2013). The diversity of the pH-reporters has also advanced through the introduction of red-shifted variants, such as mOrange2, mCherry, and mScarlett (Liu et al., 2021; Rajendran et al., 2018; Truckenbrodt et al., 2018), broadening the range of fluorescent markers available for live imaging studies. Overall, pH-based reporters offer a broader dynamic range for imaging, with excellent sensitivity, but require the exogenous introduction of genetic material and subsequent expression in neurons, which complicates experimental procedures.

A third category of vesicle-labeling tools are antibodies against the luminal domain of synaptotagmin1 (Syt1), which have been used for more than three decades (Kraszewski et al., 1995; Matteoli et al., 1992). These antibodies are added to the neurons for tens of minutes or hours, and are internalized during spontaneous exo- and endocytosis events, resulting in abundant synaptic vesicle labeling. As they specifically label synaptic vesicles, unlike the FM dyes, they do not require extensive washing steps, and do not require harsh stimulation paradigms. The latter are needed for FM dyes, since long incubations, under spontaneous culture activity conditions, would also enable the labeling of other types of vesicles and/or endosomes with the dyes, which would confuse the analyses. No such labeling is observed with the Syt1 antibodies, under normal labeling conditions (Truckenbrodt et al., 2018). However, the antibodies cannot be released by the recycling vesicles, implying that they are unable to report exocytosis. To address this problem, they have been conjugated to chemical pH-sensitive fluorophores that detect the acidic environment within vesicles, thereby mimicking the functionality of pHluorins (Hua et al., 2011). Their use remains limited, however, especially as CypHer5E, the most commonly used fluorophore, is easily bleached, and is also prone to forming aggregates that interfere with the analysis.

Nonetheless, this approach suggests an optimal use for the Syt1 antibodies. If they could be coupled to pHluorins, they would report exo- and endocytosis in the same fashion as these genetically-encoded reporters, without any need for genetic manipulations. We report here this solution, in the form of a tool comprised of an antibody against the luminal domain of Syt1, decorated with a pHluorin-conjugated secondary nanobody. A brief incubation period of these two components forms a stable complex (Pleiner et al., 2018; Sograte-Idrissi et al., 2020), allowing for easy and straightforward labeling of synaptic vesicles. This tool, termed Syt1-nb-pHluorin, integrates the advantages of both anti-Syt1 antibodies and pHluorin-tagged proteins, enabling the analysis of vesicle exocytosis, endocytosis and recycling. This approach adds a reliable, complementary assay to the existing repertoire of methods for imaging synaptic vesicle dynamics.

## Materials and Methods

### Animals

All used animals were handled according to the regulations of the University of Göttingen and of the local authorities, the State of Lower Saxony (Landesamt für Verbraucherschutz, LAVES, Braunschweig, Germany). All animal experiments were approved by the local authority, the Lower Saxony State Office for Consumer Protection and Food Safety (Niedersächsisches Landesamt für Verbraucherschutz und Lebensmittelsicherheit) and carried in accordance with the European Communities Council Directive (2010/63/EU).

### Preparation of primary rat dissociated hippocampal cultures

Newborn rats (Rattus norvegicus) were used for the preparation of dissociated primary hippocampal cultures, as previously described (Banker & Cowan, 1977; Kaech & Banker, 2006). Shortly, hippocampi of newborn rat pups (wild-type, Wistar) were dissected in Hank’s Buffered Salt Solution (HBSS, 5mM KCl, 6mM glucose, 140mM NaCl, 4mM, NaHCO3, 0.4mM KH2PO4 and 0.3mM Na2HPO4). Then the tissues were incubated for 60 min in enzyme solution (Dulbecco’s Modified Eagle Medium (DMEM, #D5671, Sigma-Aldrich, Germany), containing 0.5 mg/mL cysteine, 2.5 U/mL papain, 50mM EDTA, 100mM CaCl2, and saturated with carbogen for 10 min). Subsequently, the dissected hippocampi were incubated for 15 min in a deactivating solution (DMEM containing 0.2 mg/mL trypsin inhibitor, 0.2 mg/mL bovine serum albumin (BSA) and 5% fetal calf serum). Then the cells were triturated and seeded on circular glass coverslips with a diameter of 18mm at a density of approximately 80,000 cells per coverslip. Prior to seeding, all coverslips underwent treatment with nitric acid, sterilization, and coating overnight (ON) with 1 mg/mL poly-L-lysine. The cells were allowed to adhere to the coverslips for 1–4 h at 37 °C in plating medium (DMEM containing 3.3mM glucose, 2mM glutamine, and 10% horse serum). The plating medium was then replaced with a Neurobasal-A medium (Life Technologies, Carlsbad, CA, USA) containing 2% B27 supplement (Gibco, Thermo Fisher Scientific, USA), 1% GlutaMax (Gibco, Thermo Fisher Scientific, USA), and 0.2% penicillin/streptomycin mixture (Biozym Scientific, Germany). Before use, the cultures were maintained in a cell incubator at 37 °C, and 5% CO2 for 12–14 days. Percentages represent volume/volume.

### Production of the synaptic vesicle labeling tool

To generate our specialized synaptic vesicle labeling tool, two key components were utilized: a mouse anti-Syt1 antibody (Cat# 105 311, Synaptic Systems, Göttingen, Germany) and a custom-engineered single-domain camelid antibody (Cat# N2042, NanoTag, Göttingen, Germany). The core sequence of the single-domain camelid antibody was conjugated to the C-terminal pHluorin via a hydrophilic 47-amino acid linker, which included 9 amino acids derived from the original hinge region. This linker was followed by a spacer region containing a TEV protease recognition site (13 amino acids) and a triple FLAG tag (25 amino acids).

### Labeling

Prior labeling, the primary mouse anti-Syt1 antibody at a dilution of 1:500 and the secondary, custom made single-domain camelid antibody with concentration 1mg/ml and dilution of 1:250, were preincubated in cell culture medium (constituting 10% of the final volume for labeling) for 40 minutes at room temperature (RT). This pre-incubation step was critical for ensuring the formation of a stable complex between the primary antibody and the secondary single-domain camelid antibody. The volume of the cell culture medium solution, containing our labeling tool was then increased to the final volume needed for the labeling procedure (300 µl per coverslip). After a brief vortexing, 300 µl of labeling solution was pipetted to the wells of a new 12 well plate (Cat# 7696791, TheGeyer, Renningen, Germany). The coverslips were then transferred to the well plate. Subsequently, the neuronal cultures were incubated for 90 min at 37°C. After the incubation period, the neuronal cultures were washed 3 times in pre-heated Tyrode’s solution (containing 5 mM KCl, 30mM glucose, 2mM CaCl2, 124mM NaCl, 1mM MgCl2 and 25mM HEPES, pH 7.4) and returned to their initial well plate, containing their own conditioned media. After an additional period of incubation (60min), the cells were ready for imaging.

### Transfection Procedure

The transfection of neuronal cultures was performed using the Lipofectamine 2000 kit (#11668019, ThermoFisher Scientific, Germany) following standard protocols. Briefly, neuronal cultures were pre-incubated for 25-30 min in 400 μl of pre-heated DMEM (#D5671, Sigma-Aldrich, Germany) per well, supplemented with 10 mM MgCl_2_ at pH 7.5 (fresh-DMEM). During the incubation period, a lipofectamine mix, containing 1 μl lipofectamine solution and 24 μl Opti-MEM (#11058-021, Life Technologies Limited, United Kingdom) per well was prepared. This solution was incubated for 5 minutes at RT and added to another solution of 1 μg of the synaptophysin-mOrange2 (mOr2-sypHy) plasmid, diluted in a total volume of 25 μl Opti-MEM, per well. The final mixture was incubated for additional 15 minutes at RT and added to the neurons (in total 50 μl per well). After incubating the neurons at 37°C and 5% CO2 for 20 minutes, they were washed twice with fresh-DMEM and returned to their original culture medium. The cultures were then maintained at 37°C and 5% CO2 until the live-imaging experiments were conducted.

### Stimulation

To block recurrent neuronal activity, 50 µM AP5 (Tocris Bioscience, Bristol, UK; Abcam, Cambridge, UK) and 10 µM CNQX (Tocris Bioscience, Bristol, UK; Abcam, Cambridge, UK) and were added to the imaging solution (preheated Tyrode’s buffer). Electrical stimulation of the cells in Fig2A and B was carried with field pulses at a frequency of 20Hz, lasting for 0.4, 10, 30 or 40s at 20 mA and in Fig2C was performed at a frequency of 5 and 20Hz for 30 s with a pause of 30 s between stimulations, and followed by NH_4_CL treatment. This stimulation was performed with A310 Accupulser Stimulator and 385 Stimulus Isolator (both, World Precision Instruments, Sarasota, FL, USA), and with the help of a platinum custom-made plate field stimulator (with 8mm distance between the plates).

**Figure 1.**
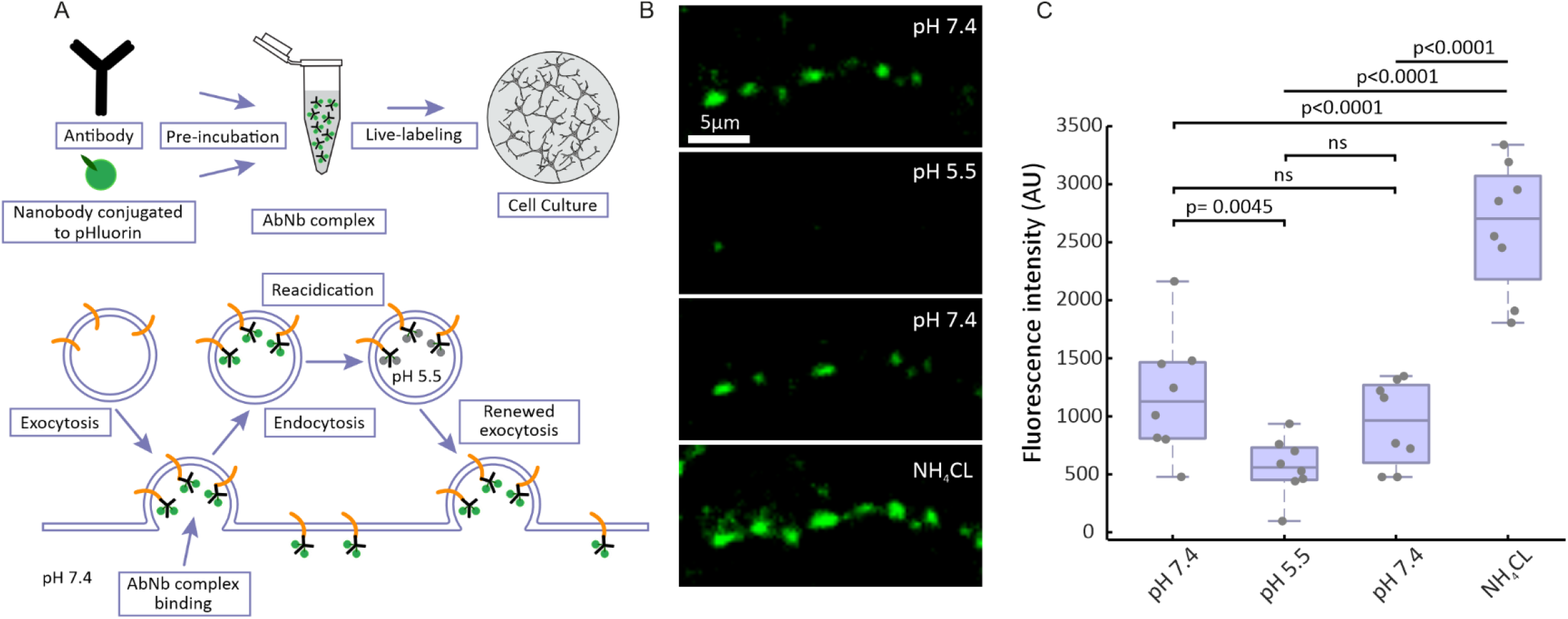
Experimental setup for recording synaptic activity. **A**. Schematic of the synaptic labeling procedure, using a primary antibody and a secondary single-domain camelid antibody (nanobody) conjugated to pHluorin. The antibody-nanobody complex is taken up by vesicles during a live-labeling period of 90 minutes, thereby labeling the entire recycling pool. The pHluorin moiety is then able to react to exo- and endocytosis, as it is quenched in the acidic interior of the synaptic vesicle, but it is allowed to fluoresce at neutral pH. **B**. Representative images of synaptic boutons subjected to different pH conditions, with 100 mM NH_4_Cl applied at the end, to induce alkalinization. **C**. Quantification of fluorescence intensity across synaptic boutons (n = 280), over 8 independent experiments (N = 8). Each black circle represents the mean value per experiment. Data are presented as a boxplot showing the median and quartiles, while whiskers represent the full data range. Statistical analysis was performed using One-Way ANOVA followed by Tukey’s LSD test.

**Figure 2.**
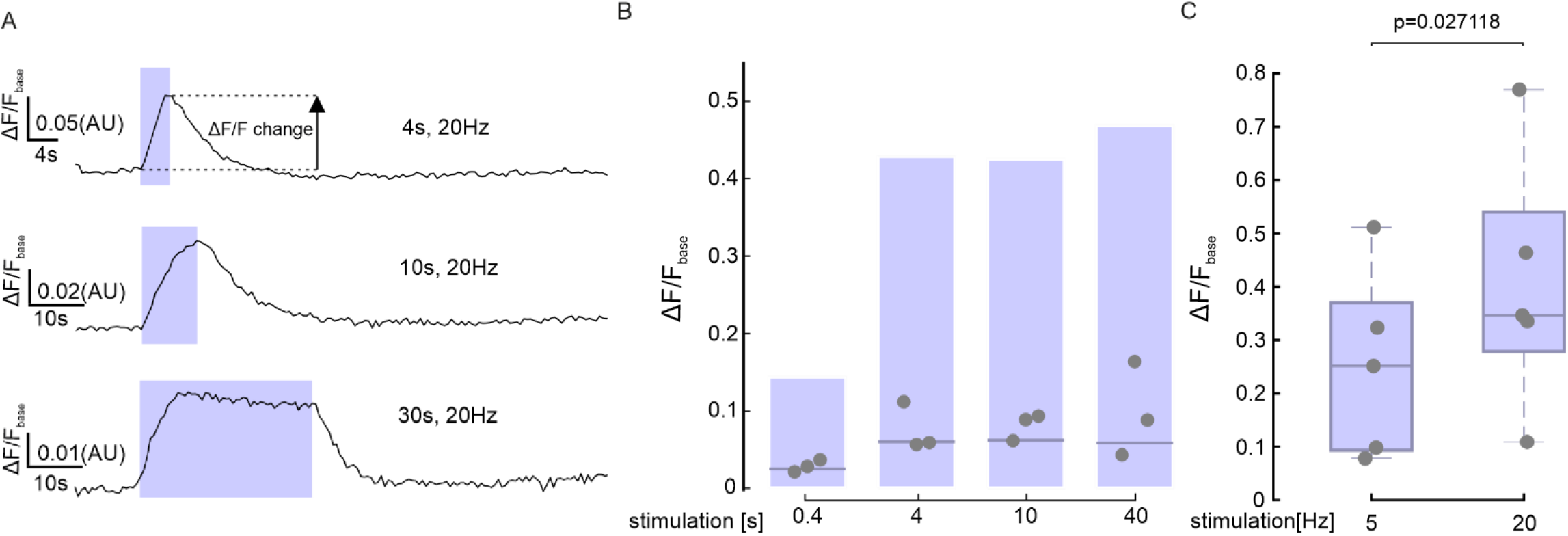
Syt1-nb-pHluorin successfully responds to various stimulation paradigms. **A**. Traces showing changes in fluorescence intensity (expressed as fluorescence normalized to the baseline signal) from 3 different experimental paradigms, in which synapses were stimulated for 4, 10 or 30 s, at 20 Hz. **B**. Flying bars showing the changes in fluorescence during the peak response (as indicated in A) to stimulations for 0.4, 4, 10 and 40 s at 20 Hz. The line indicates the median value, the edges of the box the minimum and maximum values. N=3 experiments per condition with 251, 244, 441 and 390 boutons analyzed per respective conditions. **C**. Boxplots, showing the changes in fluorescence during the peak response to stimulations at 5 Hz and 20 Hz (for 30 s), normalized to the fluorescence intensity changes after NH_4_CL treatment. N = 5 independent experiments. Statistical analysis was performed using two-tailed paired t-test.

### Live-Imaging

The coverslips were mounted on a custom-made live-imaging chamber, and 400 ul of pre-heated imaging solution was added (Tyrode’s solution supplemented with the aforementioned drugs). The neurons were then imaged with an inverted Nikon Ti microscope, equipped with Plan Apochromat 60x 1.4NA oil objective (Nikon Corporation, Chiyoda, Tokyo, Japan), a cage incubator system (OKOlab, Ottaviano, Italy), an Andor iXON 897 emCCCD Camera (Oxford Instruments, Andor), with a pixel size of 16×16μm, and Nikon D-LH Halogen 12V 100W Light Lamp House. A temperature of 37°C was maintained throughout the live-imaging procedure. The illumination and imaging frequency were as follows: 200 ms illumination and 1.7 frames per second (fps) imaging frequency for experiments in Fig2 and 200 ms illumination in both channels and 0.55 fps imaging frequency for experiments in Fig. 3.

**Figure 3.**
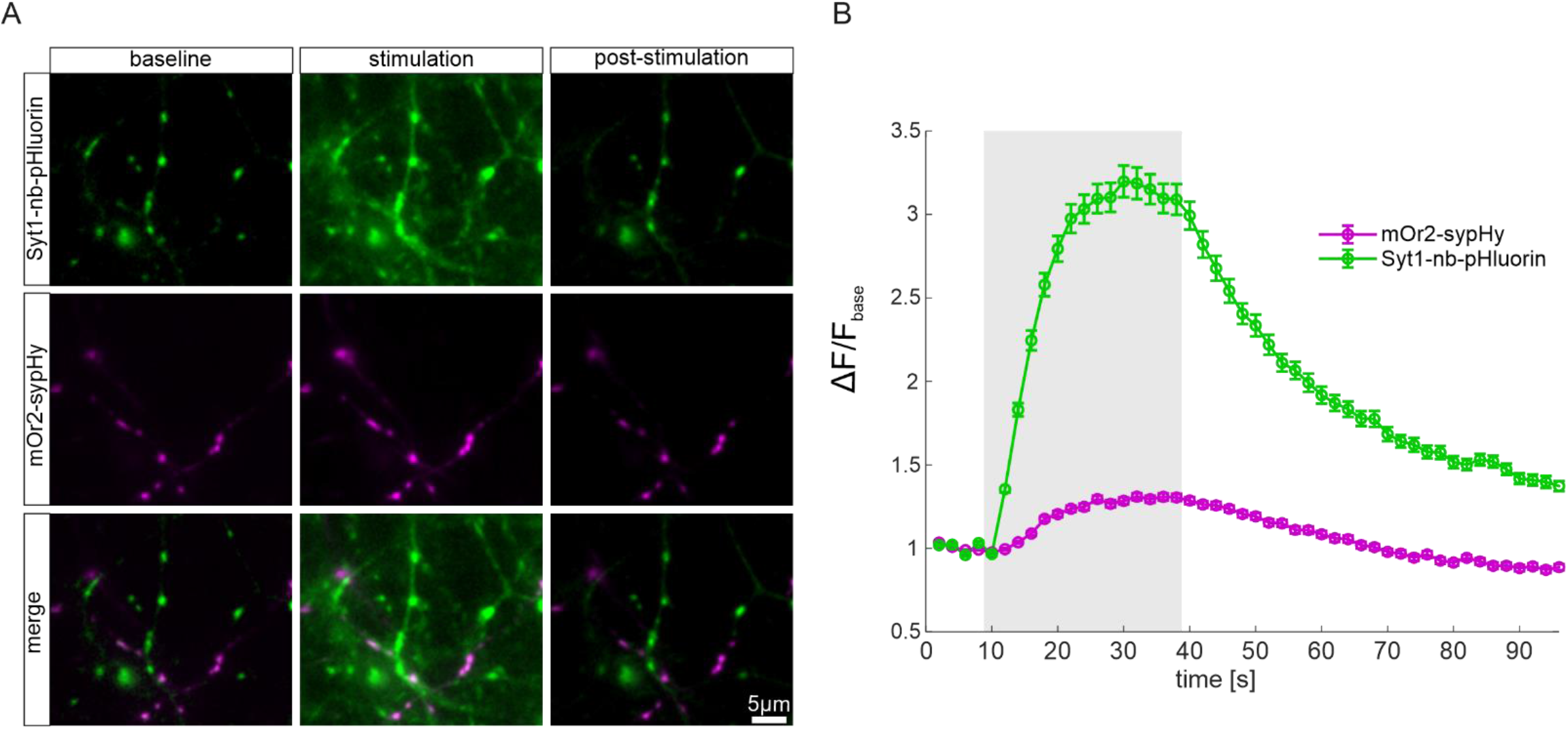
Comparison between Syt1-nb-pHluorin and Synaptophysin-mOrange2. **A**. Representative images of synaptic boutons from hippocampal neurons transfected with mOr2-sypHy and stained with Syt1-nb-pHluorin before, during and after application of a stimulus of 600 action potentials (APs) at 20 Hz. **B**. Representative signal curves from Syt1-nb-pHluorin (green) and mOr2-sypHy (magenta) in response to 600 action potentials at 20 Hz (N = 3 experiment). The error bars represent s.e.m.

### Data Analysis

The resulting movies were analyzed using custom routines developed in Matlab (The Mathworks Inc., Natick MA, USA; version R2023b). Initially, the frames were drift-corrected to account of any spatial displacement, and were subsequently summed, to generate an overall image with a signal-to-noise ratio superior to that of the individual frames. Synapse coordinates and areas were automatically identified in the summed images using a bandpass filtering procedure (Crocker & Grier, 1996), followed by thresholding with an empirically-determined threshold. For each frame, the fluorescence intensity signal within the identified synapse areas was measured, corrected for background signal, and analyzed to assess the changes induced by stimulation. These changes were quantified as the fractional change in intensity relative to the baseline.

For the vesicle pool analysis (Fig. 2C) we measured the peak signals during two rounds of stimulation, first at 5 Hz (30 seconds), and then at 20 Hz (30 seconds), and then expressed them as fraction of the respective signals obtained after NH_4_Cl dequenching. Synapses providing negative signals for any of the three measurements were removed from the analysis, as they were considered to not respond properly to stimulation.

### Statistical Analysis and Visualization

For statistical analysis Matlab (The Mathworks Inc., Natick MA, USA; version R2023b) was used. For creation of the graphs Matlab and CorelDraw (Version 24.5.0.731) were used.

## Results

### Syt1-nb-pHluorin successfully reports synaptic vesicle exocytosis

To validate the pH sensitivity of Syt1-nb-pHluorin, we conducted a classical experiment in which we manipulated the pH of the cell culture buffers and monitored the resulting changes in fluorescence intensity at synaptic boutons (Sankaranarayanan et al., 2000). We labeled neurons live by applying Syt1-nb-pHluorin to the neuronal cultures over a 90-minute incubation. The uptake of our tool occurred through spontaneous exo- and endocytosis, under conditions of spontaneous culture activity, as is typically performed for these antibodies (Truckenbrodt et al., 2018). Once endocytosed, the acidic lumen of the vesicles quenches the pH-sensitive reporter (Fig. 1A). Upon re-exocytosis, the pH-sensitive reporter is exposed to the neutral extracellular pH, allowing it to fluoresce upon illumination. As shown in Fig.1B, baseline fluorescence intensity, which reveals Syt1 molecules found on the neuronal surface (known as surface pool), significantly decreased when the extracellular pH was lowered to 5.5. Upon replacement of the acidic solution with Tyrode’s buffer at pH 7.4, the fluorescence intensity nearly returned to its initial level. We then perfused the cells with a buffer containing 100 mM NH_4_Cl. This results in the diffusion of ammonia across cell membranes, leading to the elevation of cytosolic and organelle pH (Roos & Boron, 1981). As expected, a substantial rise in fluorescence intensity was observed. These results demonstrate that Syt1-nb-pHluorin effectively labels synapses, and is capable of sensing changes in the pH at the synapse.

### Syt1-nb-pHluorin senses exo- and endocytosis upon stimulation

We next tested the performance of Syt1-nb-pHluorin under different electrical stimulation paradigms. To prevent recurrent neuronal activity, we supplemented our imaging solution with d,l-2-amino-5-phosphonovaleric acid (AP5) and 6-cyano-7-nitroquinoxa line-2,3-dione (CNQX). We employed various stimulation protocols, ranging from 8 to 800 action potentials (APs) at 20 Hz. Across all experiments, we observed significant fluorescence intensity changes relative to baseline fluorescence (Fig. 2A and B). The application of Syt1-nb-pHluorin enabled us to detect exocytosis events from the readily releasable pool of synaptic vesicles, which are immediately available for release upon short stimulations (8, 80 APs) (Fernández-Alfonso & Ryan, 2004). Additionally, we detected exocytosis from the recycling pool, which is mobilized following more sustained stimulation (200, 600, 800 APs) (Truckenbrodt et al., 2018; Wienisch & Klingauf, 2006). Moreover, our procedure enabled us to label vesicles belonging to the reserve pool, as indicated by comparing the fluorescence intensity changes from baseline after stimulations with 150 APs and 600APs, normalized to the change of fluorescence intensity signal, following NH_4_Cl application (Fig2C). The latter stimulus is suggested to release the entire recycling pool (Fernández-Alfonso & Ryan, 2004; Truckenbrodt et al., 2018; Wilhelm et al., 2010), and it appears to cause the exocytosis of only ∼30-40% of all pHluorin-labeled vesicles, which is consistent with values observed in studies investigating the reserve pool and using conventional pHluorin reporters or other tools (Fernández-Alfonso & Ryan, 2004; Truckenbrodt et al., 2018). These findings demonstrate that Syt1-nb-pHluorin is a sensitive tool that enables the analysis of exocytosis and recycling of different synaptic vesicle populations.

### Comparative analysis of Syt1-nb-pHluorin and mOr2-sypHy

To assess the efficacy of Syt1-nb-pHluorin in labeling synaptic boutons and studying vesicle recycling, relative to existing methods, we conducted dual-color live imaging experiments on hippocampal neuronal cultures transfected with synaptophysin-mOrange2 (mOr2-sypHy). The mOr2-sypHy construct is a modified version of the original sypHy probe, wherein the superecliptic GFP is replaced by the pH-sensitive mOrange2 fluorescent protein (Shaner et al., 2008; Truckenbrodt et al., 2018). Due to the transient plasmid expression used for mOr2-sypHy, we observed relatively low transfection efficiency, resulting in only a few labeled cells per coverslip. In contrast, our Syt1-nb-pHluorin tool demonstrated substantially higher labeling efficiency across the neuronal cultures, as it is expected to label all active synapses. Nevertheless, we successfully labeled functional synaptic boutons expressing mOr2-sypHy with Syt1-nb-pHluorin, as illustrated in Figure 3A. We quantified the fluorescence intensity changes relative to baseline (ΔF/F_ase_) in these dual-labeled boutons, upon stimulation with 600 APs. As depicted in Figure 3B, Syt1-nb-pHluorin exhibited a significantly greater dynamic range, when compared to mOr2-sypHy, indicating a more robust detection of synaptic vesicle exocytosis events. This is in line with previous results (Truckenbrodt et al., 2018), which indicate that the pH-dependent changes in mOr2 fluorescence are lower than those undertaken by the GFP-based pHluorin. Nonetheless, the kinetics observed with the two fluorophores are similar, indicating that the Syt1-nb-pHluorin provides qualitatively identical results to genetically-encoded detectors.

## Discussion

In this study, we demonstrated that a tool composed of primary antibodies against Syt1 and secondary nanobodies conjugated to pHluorin (Syt1-nb-pHluorin) exhibits robust performance in monitoring synaptic vesicle dynamics in primary hippocampal rat neuronal cultures. It labels synaptic boutons and detects pH-changes in their vesicles, characteristics that render it a reliable pH reporter. Syt1-nb-pHluorin also responds to a wide range of stimulation paradigms, from 8 APs to 800 APs, indicating its ability to track exocytosis across different synaptic vesicle pools, including the readily releasable pool or recycling vesicles. Syt1-nb-pHluorin offers superior labeling efficiency improved signal to noise ratio, compared to reporters as mOrange2. These features make Syt1-nb-pHluorin a convenient tool for studying synaptic vesicle recycling.

While our tool shows promising performance in studying synaptic vesicle dynamics, several challenges remain to be addressed. Although it avoids the need of extensive washing steps, as in the case of FM dyes, it still requires a pre-incubation period, during which the antibody-nanobody complex is formed and then taken up by the vesicles. This step typically takes approximately 130 minutes, and requires primary antibody to secondary nanobody ratios of at least 1:2, to ensure that all binding sites on the primary antibody are occupied by the pH-conjugated nanobody (Sograte-Idrissi et al., 2020). Nevertheless, this process is faster than conventional transfection techniques, as lipofectamine or calcium phosphate transfection, and, crucially, it also bypasses the need for genetic manipulation of neurons. A possible solution to this challenge would be the direct conjugation of the primary antibody to a pH-reporter. This has already been employed with tools like CypHer5E fused to Syt1(Hua et al., 2011), a pH reporter that operates in an opposite manner to most pHluorins, since it is quenched at neutral pH, and is activated by the acidic environment inside the vesicles. A promising next step would involve the development of nanobodies against Syt1, that are directly conjugated to pHluorin. Syt1 nanobodies against the cytosolic parts of Syt1 already exist, and have been successfully employed in neurophysiological studies (Queiroz Zetune Villa Real et al., 2023), but nanobodies against its luminal domain still need to be characterized.

Another significant challenge when using Syt1-nb-pHluorin is the limited fraction of synaptic vesicles labeled during the pre-incubation procedure. This limitation arises from the fact that Syt1-nb-pHluorin specifically labels the luminal epitopes of Syt1 that are exposed to the extracellular milieu during spontaneous exo- and endocytosed events, thereby exclusively targeting vesicles that undergo active exo- and endocytosis. The reserve pool of vesicles, which remains inactive during physiological conditions, only fuses with the plasma membrane following prolonged, non-physiological electrical stimulations, or the application of pharmacological agents of drugs that enhance presynaptic activity (Frontali et al., 1976; Kim & Ryan, 2010; Rizzoli & Betz, 2005). To label reserve vesicles, the user needs to wait a significant time period after labeling (preferably hours), to ensure that the newly recycled vesicles intermix functionally with the reserve pool (Truckenbrodt et al., 2018). Nevertheless, the incubation time of 60 minutes, after the 90-minutes labeling procedure, was sufficient to label at least a fraction of the reserve vesicle pool, in our experiments (Fig2C).

In conclusion, we believe that our tool is a valuable contribution to the existing methods for studying synaptic vesicle dynamics.

## Acknoledgements

We thank Dr. Eugenio Fornasiero and Ronja Rehm (University Medical Center Göttingen, Germany) for support with imaging-related procedures. We thank Christina Zeising and Nicole Hartlet (University Medical Center Göttingen, Germany) for expert technical assistance. We thank Dr. Guobin Bao for the help during the stimulation-related procedures. S.O. Rizzoli and S.V. Georgiev were funded by the German Research Foundation (Deutsche Forschungsgemeinschaft, DFG), through the Collaborative Research Center (CRC) 1286, project A03. S.O. Rizzoli and S.V. Georgiev were further supported by the Deutsche Forschungsgemeinschaft under Germany’s Excellence Strategy–EXC 2067/1-390729940.

## Notes

### Competing Interest Statement

The authors have declared no competing interest.

